# The 3-D structure of fibrous material is fully restorable from its X-ray diffraction pattern

**DOI:** 10.1101/2021.02.09.430379

**Authors:** Hiroyuki Iwamoto

## Abstract

X-ray fiber diffraction is potentially a powerful technique to study the structure of fibrous materials, such as DNA and synthetic polymers. However, only rotationally averaged diffraction patterns can be recorded, and it is difficult to correctly interpret them without the knowledge of esoteric diffraction theories. Here we demonstrate that, in principle, the non-rotationally averaged 3-D structure of the material can be restored from its fiber diffraction pattern. The method is a simple puzzle-solving process, and in ideal cases, it does not require any prior knowledge about the structure, such as helical symmetry. We believe that the proposed method has a potential to transform the fiber diffraction to a 3-D imaging technique, and will be useful for a wide field of life and materials sciences.

## 1. Introduction

Like electron microscopy, X-ray diffraction technique (XRD) is a powerful tool for atomic-resolution structure analysis. However, structure analysis by XRD is more complicated than visual observation by electron microscopy, because X-ray detectors cannot record the phase of scattered waves. Reconstruction of high-resolution images requires both phase and amplitude information of scattered waves. Because of this, the structure analysis by XRD is almost synonymous to the retrieval of the once-lost phase information. Once it is done, XRD functions as X-ray microscopy with molecular-to-atomic resolution.

Fortunately, phase retrieval methods have well been established in X-ray crystallography (Hauptman, 1986; Taylor, 2010). Even for non-crystalline XRD, Algorithms has been developed to determine the structures of proteins dissolved in solutions (Svergun & Stuhrmann, 1991; Chacon et al., 1998, Grant, 2018) and isolated single particles (coherent diffractive imaging, or CDI) (Miao & Sayre, 2000). However, X-ray fiber diffraction remains as the last frontier of non-crystalline XRD with no established phase retrieval methods.

Classically, X-ray fiber diffraction contributed to the discovery of the double-helical structure of DNA (Watson & Crick, 1953), and today it is applied to a very wide range of materials, from biomolecules to synthetic polymers. Modern synchrotron-based fiber diffraction is the only means that can monitor sub-millisecond molecular motions within functioning fibers, e.g., in the flight muscle of a flying insect (Iwamoto & Yagi, 2013).

In the absence of good phase-retrieval methods, however, information extractable from fiber diffraction is limited. A further problem is that one can usually record only rotationally averaged diffraction patterns, from which angular information is missing. Attempts have been made to apply the CDI technique to fibrous materials (Latychevskaia & Fink, 2018), but it is not well suited for fibrous materials, because continuous scattering densities required for CDI (Miao & Sayre, 2000) is not usually obtained.

Here we demonstrate that, in principle, the non-rotationally averaged 3-D structure of a fiber can be restorable from a single rotationally averaged diffraction pattern, without prior knowledge about the symmetry of fiber structure. This is based on the Patterson method (Patterson, 1934), one of the direct phase-retrieval methods developed in a very early stage of the X-ray diffraction science. The method is based on a simple and clear procedure, and in most cases, the 3-D structure is uniquely determined. Unlike in CDI, no iterative calculations are required.

## 2. Principle

When both amplitude and phase are available, the inverse Fourier transformation (F^-1^) of the scattered waves generates the image of the sample at a high resolution, and this is actually the principle of lens-based microscopes. If one performs Fourier transformation on the intensity of diffraction pattern, one obtains a Patterson function instead. This is an auto-correlation function of the sample structure, and is the list of all the vectors that connect any pairs of masses within the sample. A Patterson function can be calculated from a diffraction pattern and, if the sample structure is known, it can also be calculated from it. The Patterson function for a 3-D sample is also a 3-D function.

In the case of fiber diffraction, the vectors are rotated around the fiber axis to fall on a single plane, so that the Patterson function is reduced to 2-D (the angular information around the fiber axis is lost). This is called the cylindrically averaged Patterson function (CAP) (MacGillavry & Bruins, 1948). CAP has been utilized in the analyses of fiber diffraction patterns from DNA (Franklin & Gosling, 1953), muscle (Namba et al., 1980; Ohshima et al., 2011), etc, and it gives some insights into structural symmetry, but its true potential has not been exploited.

CAP’s true potential is, as we propose here, that the non-rotationally-averaged 3-D structure of the sample can be restored from it. The vectors listed in the CAP are like the piled pieces of a zig-saw puzzle, and our mission is to tile (re-connect) them in the 3-D space to finish the puzzle. The rules are simple: (1) the vectors can only be translated, rotated around the fiber (z-) axis and inverted with respect to the x-y plane. (2) No vectors in the CAP must be left unused. (3) One may not create any vector that is not listed in the CAP. (4) A single vector may be used many times, if necessary.

We developed a computer program that actually solve the puzzles by following the rules listed above. Detailed information about the program is described in the Supplemental Material, and the results of application to model structures are described below.

## 3. Examples of implementation using model structures

The simplest example is shown in Fig. 1. Here the sample consists of only 3 atoms (a, b and c), positioned at the vertices of an equilateral triangle (Fig. 1A). The triangle is freely rotatable around the z-axis, so that its diffraction pattern is rotationally averaged (Fig. 1B). From this diffraction pattern one obtains the CAP (Fig. 1C). A CAP is symmetrical with respect to the x- and z-axes, so that the information in just one quadrant is sufficient. Here are 3 vectors in each quadrant, corresponding to vectors ab, ac and bc in the triangle. Now we start to reconstruct the triangle (Fig. 1E). First, we choose one of the vectors (here, ac) as a starting vector, and place it on the x-z plane. Next, we try to connect vector bc to vector ac. Vector bc may start from either end of vector ac, and here we tentatively connect vector bc to the origin, and place it on the x-z plane, too. Note here, that placing a new vertex in the 3-D space creates new vectors (labeled new in Fig. 1D) connecting it to all the vertices that are already present. This vector is not listed in the CAP, so the rule 3 (see above) is violated. If, however, vector bc is rotated around the z axis, at an angle, the new vector coincides with vector ab. Now all the vectors in the CAP are used and the non-rotationally-averaged 3-D structure of the triangle is successfully restored (Fig. 1F, see also supplementary movie).

**Fig. 1.**
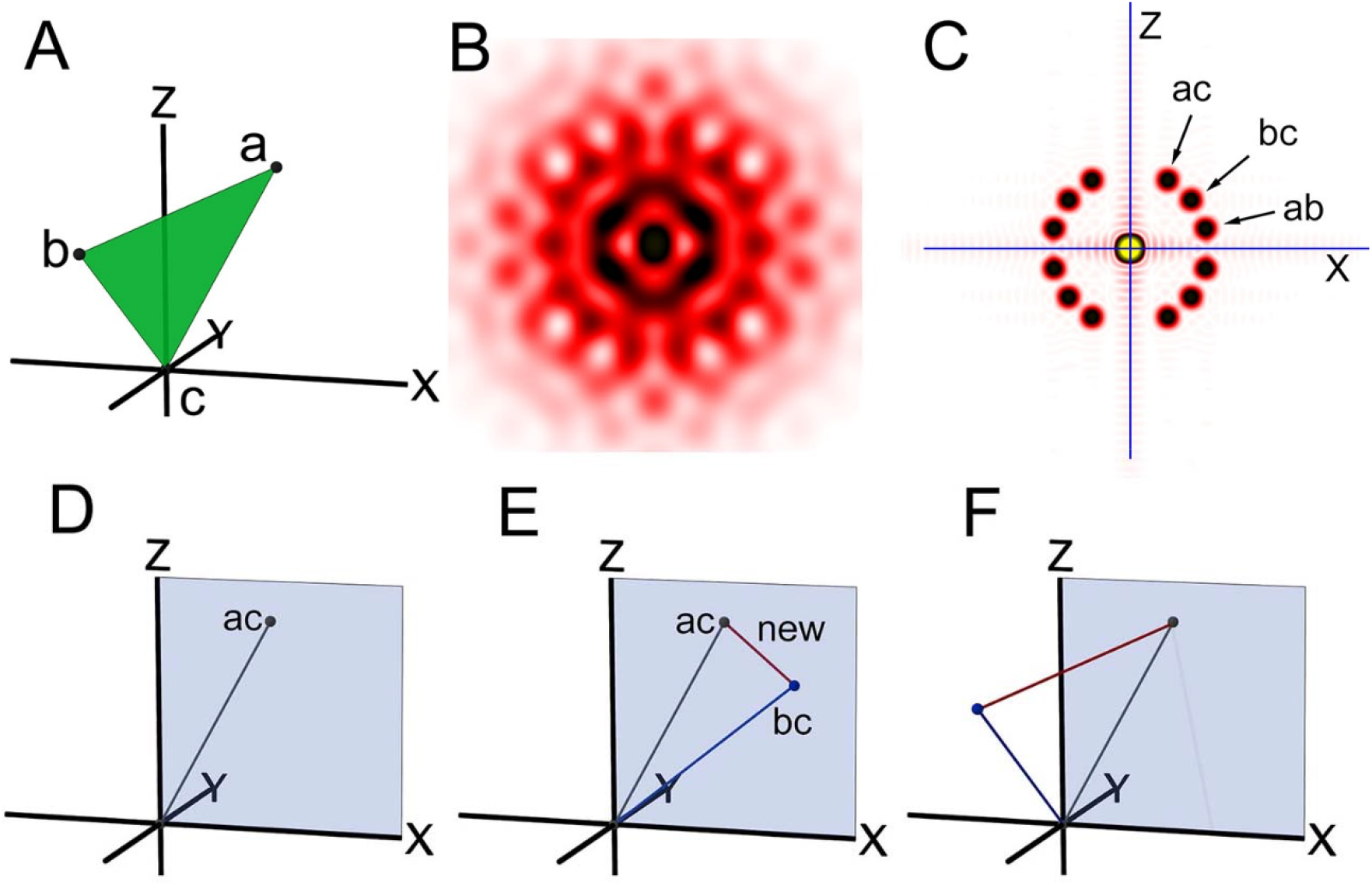
Reconstruction procedure of a 3-D structure from its cylindrically averaged Patterson function (CAP). (A) The structure of an object, consisting of only 3 points (The green plane is for assisting purpose only), freely rotatable around the z-axis. (B) “Fiber” diffraction pattern from the object. (C), CAP of the object, calculated from (B). (D)-(F) Reconstruction procedure. (D) One of the vectors (here vector ac) is placed as the starting vector on the x-z plane (pale blue), (E) A second vector (bc) is tentatively placed on the x-z plane and connected to the first vector, and this creates a new, incorrect vector (red). (F) The second vector is rotated around the z-axis, and now the new vector coincides with the unused vector ab, and the reconstruction is finished.

This is a very simple puzzle game, but it works for more complex structures. Figure 2 shows examples of the double-helix of DNA and the helix of actin filament (28 monomers in 13 turns, or 28/13 helix). For these structures, the CAPs are much more complex, but the computer program quickly finds right answers, as long as an ideal CAP is available (i.e., all the vectors are listed and their values are accurate).

**Fig. 2.**
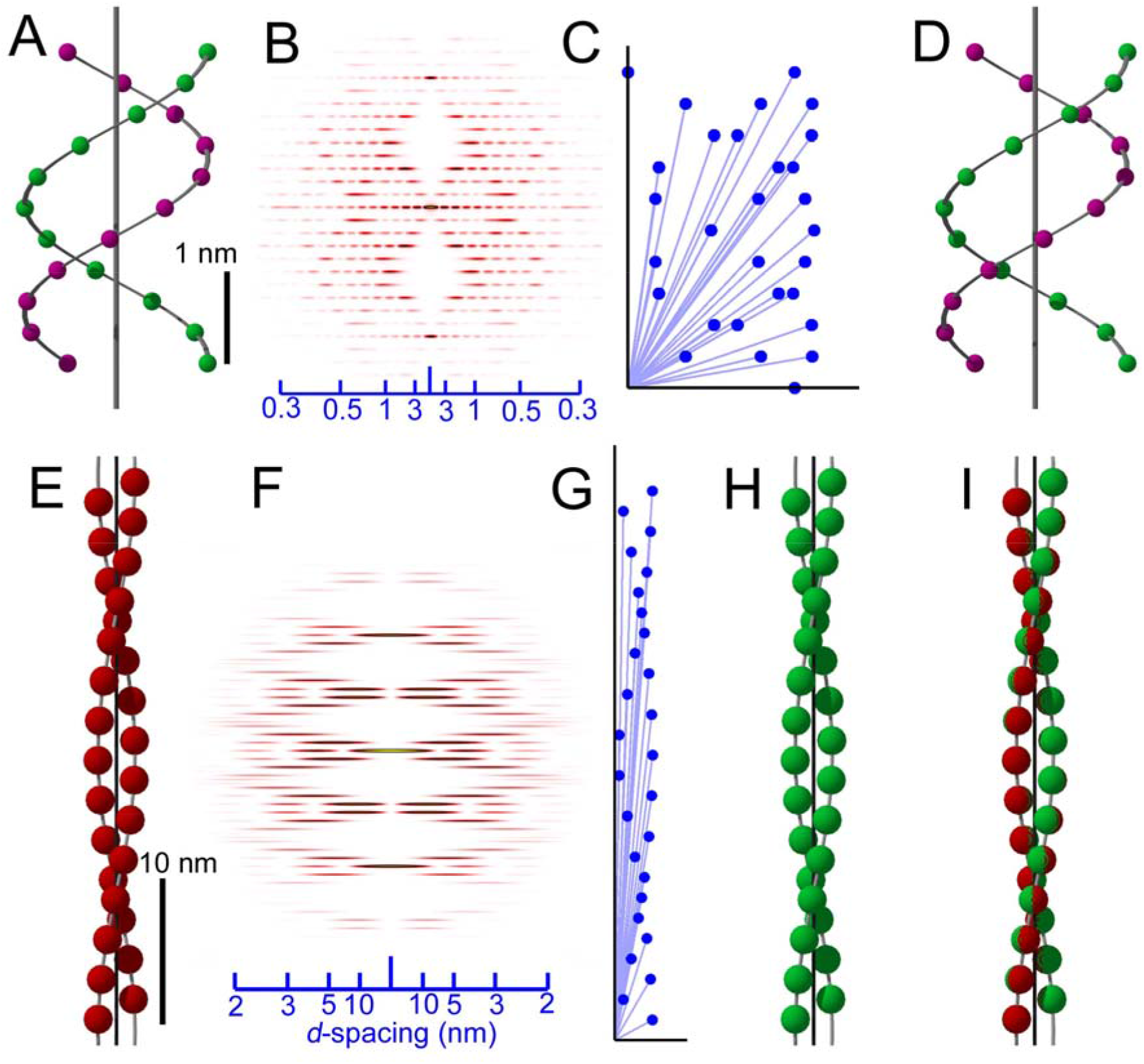
Application of the reconstruction method to helical structures. (A) and (E), the starting model structures of double-stranded DNA (phosphate positions only) and the 28/13 helix of F-actin, respectively. The gray continuous helices and the central axes are shown for assisting purposes. (B) and (F), expected fiber diffraction patterns, consisting of helix-derived layer line reflections. (C) and (G), CAPs of the model structures. (D) and (H), the 3-D structures reconstructed from the CAPs alone. (I), goodness of fit of the original and reconstructed F-actin structures, which depends on the magnitude of permissible errors set in the computer program. The scale bar in (A) also applies to (C, D), and that in (E) also applies to (G, H, I). The values on scale bars in (B, C) are *d*-spacings (nm). Supplementary Figs. 2 and 3 show how internal vectors in the 3-D structures (A, E) are picked up in their CAPS (C, G).

The handedness of the helix cannot be determined by diffraction data alone, because the diffraction patterns from opposite-handed helices are identical. For this reason

## 4. Application to an actual diffraction pattern

In actual recordings, the CAPs calculated from diffraction patterns are not usually ideal for many reasons: fused peaks, missing peaks, false peaks, inaccurate peak positions, etc. Even in such difficult cases, some prior knowledge about the symmetry of the structure can help restore the 3-D structure from the CAPs. Figure 3 shows an example: the myosin filament of insect (bumblebee) flight muscle.

**Fig. 3.**
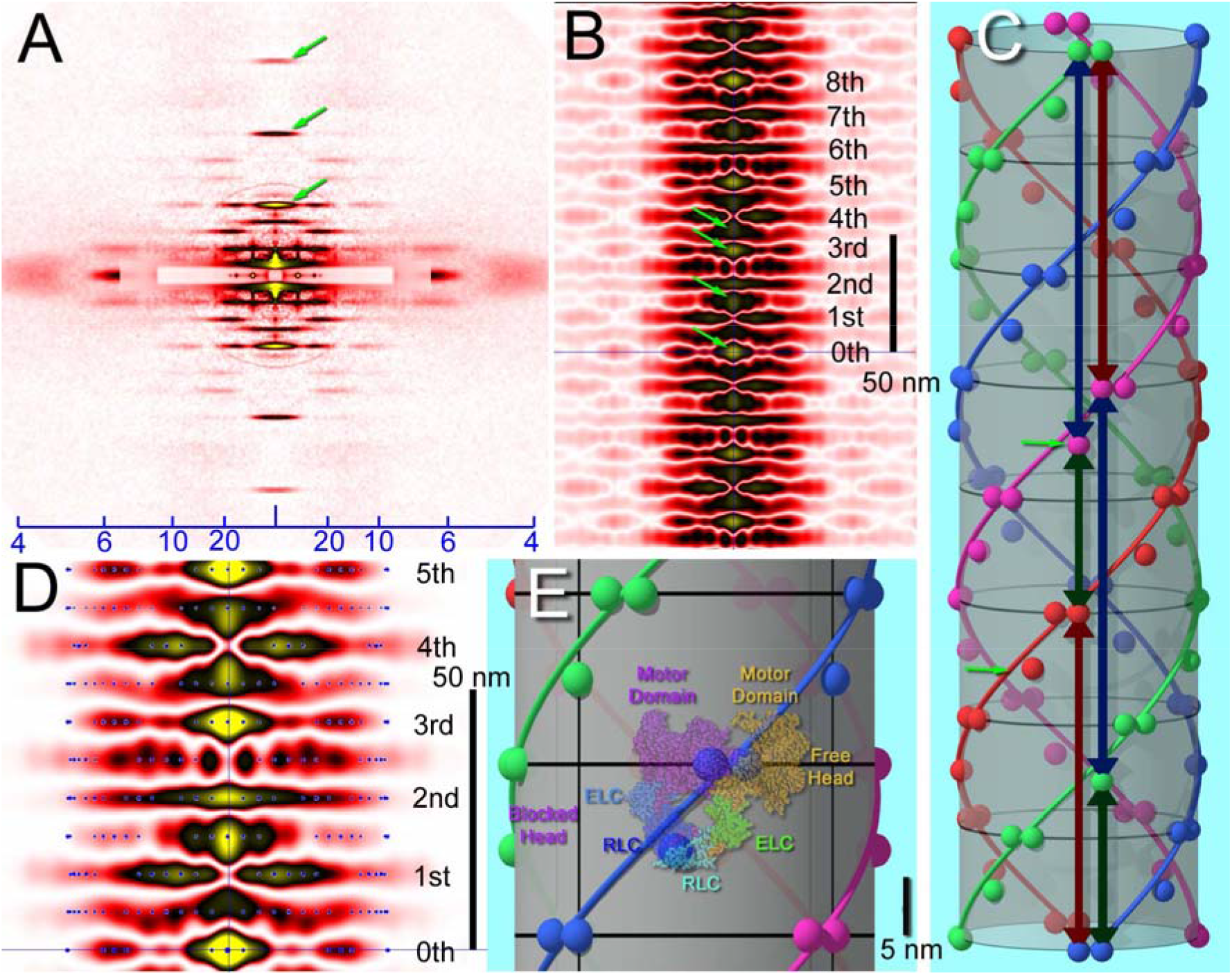
Reconstruction of the 3-D structure of myosin filament from an actually recorded diffraction pattern. (A) Diffraction pattern from actin-extracted bumblebee flight muscle fibers. Arrows indicate meridional reflections from myosin-head crowns. (B) CAP calculated from the diffraction pattern in (A). The numbers indicate the levels of the 14.5-nm crowns of myosin heads. Arrows indicate the peaks on the z-axis at the 1.5th, 3rd and 3.5th levels of myosin crowns. (C) The solved structure of the myosin filament. The black lines on the cylinder indicate the levels of crowns, and the small arrows indicate the additional mass lying midway between two neighboring crowns. The green, red and blue double-headed arrows indicate the vertical vectors of 1.5, 3 and 3.5 x 14.5 nm-lengths, respectively. (D) Superposition of the actually observed CAP and the idealized CAP calculated from the solved structure (blue dots). (E) Superposition of the interacting heads motif (Sweeney, 2018) and the three-mass unit on the solved structure. The size of the motif was adjusted by using the scale bar in (Padron, 2016) (right). The values on scale bars in (A) are *d*-spacings (nm).

Figure 3A shows the small-angle diffraction pattern from demembranated, actin-extracted bumblebee (*Bombus*) flight muscle fibers, containing myosin filaments alone (see Supplemental Materials for the methods of preparation and recording). Like DNA and actin strands, myosin filaments have a helical structure. As a result, its diffraction pattern also consists of many layer-line reflections, as in Fig. 2B and F. A few meridional reflections are also observed, and they originate from the 14.5-nm axial repeat of myosin-head crowns.

The CAP calculated from the diffraction pattern (Fig. 3B) has many fused peaks. It consists of a stack of densities that occur every 14.5nm, but densities also occur midway between the two 14.5-nm layers, indicating the presence of mass in the middle of the crowns. In this CAP, the profile at the 8th level of myosin crown is very similar to that at the 0th, and the entire pattern is symmetrical with respect to the 4th level. This feature is consistent with the 4-start, 8/1.25 helical symmetry of myosin heads in the flight muscle of giant waterbug (*Lethocerus*) (Tregear et al., 2004). Therefore, the two insects seem to share the same helical symmetry. Further observation of the CAP shows that there are peaks on the z-axis at 1.5th, 3rd and 3.5th levels, indicating the presence of vertical vectors with lengths 1.5, 3 and 3.5 x 14.5 nm. From these features alone, the filament structure is solved without the need of knowing all the vectors, as is shown in Fig. 3C. At each crown, two masses lie horizontally, separated by 11.25 degrees. In the midway between the crowns, there is an additional mass, slightly offset from the helical path of the masses on the crown.

From this solved structure, an idealized CAP can be calculated and it is overlaid on the observed CAP at a higher magnification in Fig. 3D. Its peak positions (blue dots) generally show good match with the observed densities. Manual fitting of the two CAPs indicates that the radius of the myosin heads from the center is ~15.3 nm.

Overall, the arrangement of the three masses in the structure in Fig. 3C is reminiscent of the interacting heads motif, in which one of the two heads of a myosin molecule blocks the other head (Zhao et al., 2009). This motif is commonly observed in many animals (Alamo et al., 2018). In Fig. 3E, the atomic model of the motif (Sweeney, 2018) is superposed on the solved structure. If this superposition is correct, the two masses on the crown correspond to two motor domains, and the mass between the crowns, the light-chain domains. In *Lethocerus* flight muscle, all the domains are at the crown level (Hu et al., 2016), and this may reflect an inter-species difference.

## 5. Future prospects and conclusion

This work demonstrates that the non-rotationally averaged 3-D structure of a fibrous material is restorable from its diffraction pattern by using CAP. Under ideal conditions, no prior knowledge is required. If reflections extend to an atomic range (a *d*-spacing of 0.3 nm or better), the 3-D structure with an atomic resolution will be obtained.

It should be made clear that this work presents the principle of the method, and deals with the situation in which a CAP correctly list up the internal vectors of the sample. However, CAPs calculated from actual diffraction patterns deviate from it for various reasons besides those listed in section 4, e.g., the curvature of the Ewald sphere and the distortion resulting from projection to flat detectors. In small-angle recordings these effects may be ignored, but in wide-angle recordings, these effects are substantial and the data should be corrected before the present method is applied.

The interference between neighboring helices is another issue. Ideally the helices in the sample should be infinitely diluted (as in protein solution scattering experiments), but for fibrous material it is difficult to do so. In some materials the helices are organized into a lattice, and in this case, inter-helical vectors appear in the CAP in addition to intra-helical vectors. However, inter-helical vectors appear outside the intra-helical vectors (in Fig. 3B, the weaker densities on both sides are considered to be inter-helical). One can simply exclude inter-helical vectors from calculations to obtain the structure of a single helix.

One may think that cryoelectron microscopy (cryo-EM) is better suited for high-resolution 3-D structure determination than XRD. In principle, however, one should be able to determine atomic-resolution 3-D structure by using this protocol if one can record diffraction data up to atomic-resolution q-ranges. As briefly stated in Introduction, XRD is good at dynamic measurements, and does not usually requires pre-treatments that may affect the native structure, while in cryo-EM the sample must be cryofixed and is only capable of static measurements. In cryo-EM, a huge number of micrographs must be collected and class-averaged, but in XRD, especially that using synchrotron radiation, a single-shot picture often provide sufficient data quality. Thus, high-throughput data acquisition is possible in XRD. Then the present method will provide a quick means for 3-D structure assessment on site.

The basic method shown here may be further developed by utilizing unused features of CAP, e.g., the spread of peaks may be used to volume-render the restored structure. The imperfectness of real data may be addressed by machine-learning-based inference. It is expected that the publication of this new method stimulates interested scientists (especially data and computer scientists) to join and accelerate further development of the method. In the end, the X-ray fiber diffraction will be an easier-to-use tool to visualize individual atoms or molecules in synthetic and naturally-occurring fibers.

## Acknowledgment

We thank Drs. T. Hikima and T. Hoshino, RIKEN, for their help at the beamlines. The X-ray diffraction studies were done under approval of SPring-8 proposal review committee (2014B1260, 2015B1420, 2016A1220, 2018A1305, 2018B1431, 2019B1400). Funding: This work was supported by JSPS KAKENHI Grant No. 19K06777.

## Supplemental document

### Further description of structure reconstruction procedure

#### 1. Basic idea

A CAP can be calculated from the structure of a sample consisting of a finite number of vertices distributed in a 3-D space. When a sample consists of n vertices, the number of independent vectors nv in the CAP (those in the 1st quadrant) is n(n – 1)/2. This does not include zero-length vectors. Either in the Cartesian or cylindrical coordinates, each vertex carries 3 variables to be determined, so that the total number of variables to be determined is 3(n – 1). On the other hand, each vector in the CAP is 2-dimensional, so that it provides 2 equations. Therefore, the total number of the simultaneous equations is n(n – 1). When n ≧3, n(n – 1) is always equal to or greater than 3(n – 1), so that the simultaneous equations should be solvable and the phase information should be restorable.

If all of the independent vectors are known, the number of vertices n is calculated by using the following equation:

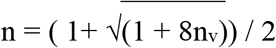

The program determines the x, y and z coordinates of all the vertices in the sample, by using a set of nv vectors. When the coordinates of all the expected n vertices have been determined, the calculations can be terminated.

When a CAP is calculated from the diffraction pattern, the exact number of vectors is usually unknown, and it is assumed that there are a number of overlapping vectors (i.e., the x and z coordinates on the CAP are identical but connecting different sets of vertices). In this case, attempts to find new vertices are continued even after all the vectors are used. When no more vertices are found and all the vectors have been used, the reconstructed 3-D structure is regarded correct (a 3-D structure consistent with the CAP has been generated).

#### 2. Specific steps

The specific steps performed in the program are described below. The software will be made available in an institutional server. The steps are visualized in the form of a flow chart in Supplementary Fig. 1.

1. Read a CAP file, which is simply a list of the (x,z) coordinates of the tips of vectors. All the vectors should be in the 1st quadrant. If a vector is identical to one of the already registered vectors, it is not registered.
2. Pick up one of the registered vectors, and place it on the x-z plane of the 3-D space as the first vector. One end of the vector is placed on the origin.
3. The other end of this vector is defined as the second vertex. The two vertices are regarded correct, and the coordinates of the rest of the vertices are determined relative to them. After this, the rest of the vertices are determined by repeating steps (4)-(11).
4. Choose two vertices that already exist, and to each of them, connect a registered vector.
5. Check if the z-coordinates of the free ends of the two connected vectors coincide.
6. If the answer for (5) is yes, further check if the two free ends can be connected. This can be checked if the circular paths along which the free ends can move cross each other.
7. If the answer for (6) is yes (There are usually two ways of connection. Both will be tested for their validity in the following steps), then the two vectors are connected, and the connection point is tentatively assigned as a new vertex.
8. When a new vertex is tentatively assigned, it creates vectors with all other existing vertices. It should be verified that all of them are those listed in the CAP. If it has been verified, the new vertex is regarded correct and the process proceed to step (9). If the new vertex creates a vector that is not in a list, the new vertex is regarded incorrect, and the process goes back to (4) and tries another combination of vectors.
9. The new vertex if formally regarded as a correct vertex.
10. Check if all the registered vectors has been used. If not, the process goes back to (4) to find more vertices, at least until no unused vector is left. If it is impossible to eliminate unused vectors, it is likely that an incorrect structure has been reconstructed. In that case, calculations are restarted from step (2) under different conditions, e.g., by using an unused vector as the starting vector.
11. Even if all the registered vectors have been used, it is checked if there are any undiscovered vertices that can be placed without creating unlisted vectors. This is done by going back to the step (4).

If no more new vertices are found, the calculation is finished, and the reconstructed 3-D structure is regarded correct (consistent with the CAP).

### Materials and methods of diffraction recordings from muscle fibers

#### 1. X-ray diffraction recording

Bumblebees (*Bombus ignitus*) were collected at or near the campus of SPring-8, and their thoraces were half-split and the flight muscle fibers in them were demembranated in a 50% mixture of a relaxing solution and glycerol, as described (Iwamoto, 2017). The fibers were stored in a 25%:75% mixture of the relaxing solution and glycerol in a −80°C freezer until use. The actin filaments within the flight muscle fibers were removed by treating the fibers with gelsolin (Kawai & Ishiwata, 2006). Bovine gelsolin (Sigma-Aldrich) was dissolved in 2 mM CaCl_2_ at a 2 mg/ml concentration, and the buffer was exchanged to an extraction buffer (80 mM K-propionate, 10 mM EGTA, 10.4 mM CaCl_2_, 5 mM MgCl_2_, 12 mM ATP, 100 mM butanedione monoxime, 20 mM imidazole, pH = 7.2) by using a NAP-5 column (GE healthcare life sciences). 5-drop fractions were collected, and the most concentrated fraction was used for extraction. The extraction was done overnight in a refrigerator, and the solution was then exchanged to a normal relaxing solution before the diffraction experiments. The extent of removal of actin filament was checked by observing the intensities of the actin-based layer line reflections in the X-ray diffraction patterns, as well as by gel electrophoretic patterns. Static X-ray diffraction patterns were recorded in the relaxing solution, in the BL45XU or BL05XU beamlines of SPring-8, as described (Iwamoto, 2009).

#### 2. Processing of X-ray diffraction data

The recorded diffraction patterns were summed and the 4 quadrants were folded to improve the signal-to-noise ratio, and the background scattering was subtracted, as described (Iwamoto et al., 2003). Then the vertical widths of the reflections of interest were reduced to 1 pixel by calculating their integrated intensities along the equator. Other reflections were deleted. This fine-lined diffraction pattern was then subjected to Fourier transformation to obtain the cylindrically averaged Patterson function (CAP). The intensities on the equator were not used, so that the obtained CAP corresponds to the difference cylindrically averaged Patterson function (Namba et al., 1980).

### Supplementary Figures

**Supplementary Fig. 1.**
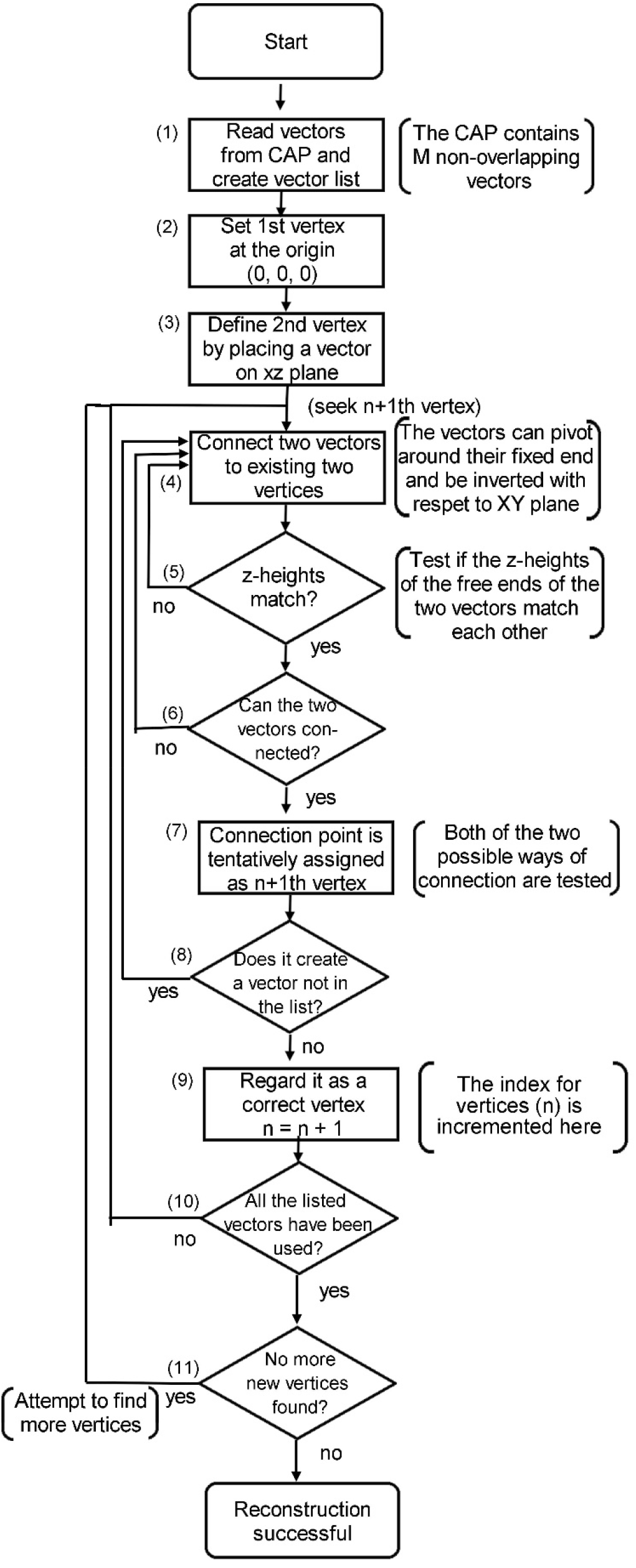
Flow-chart of the reconstruction procedure of 3-D structure from CAP. For details see text.

**Supplementary Fig. 2.**
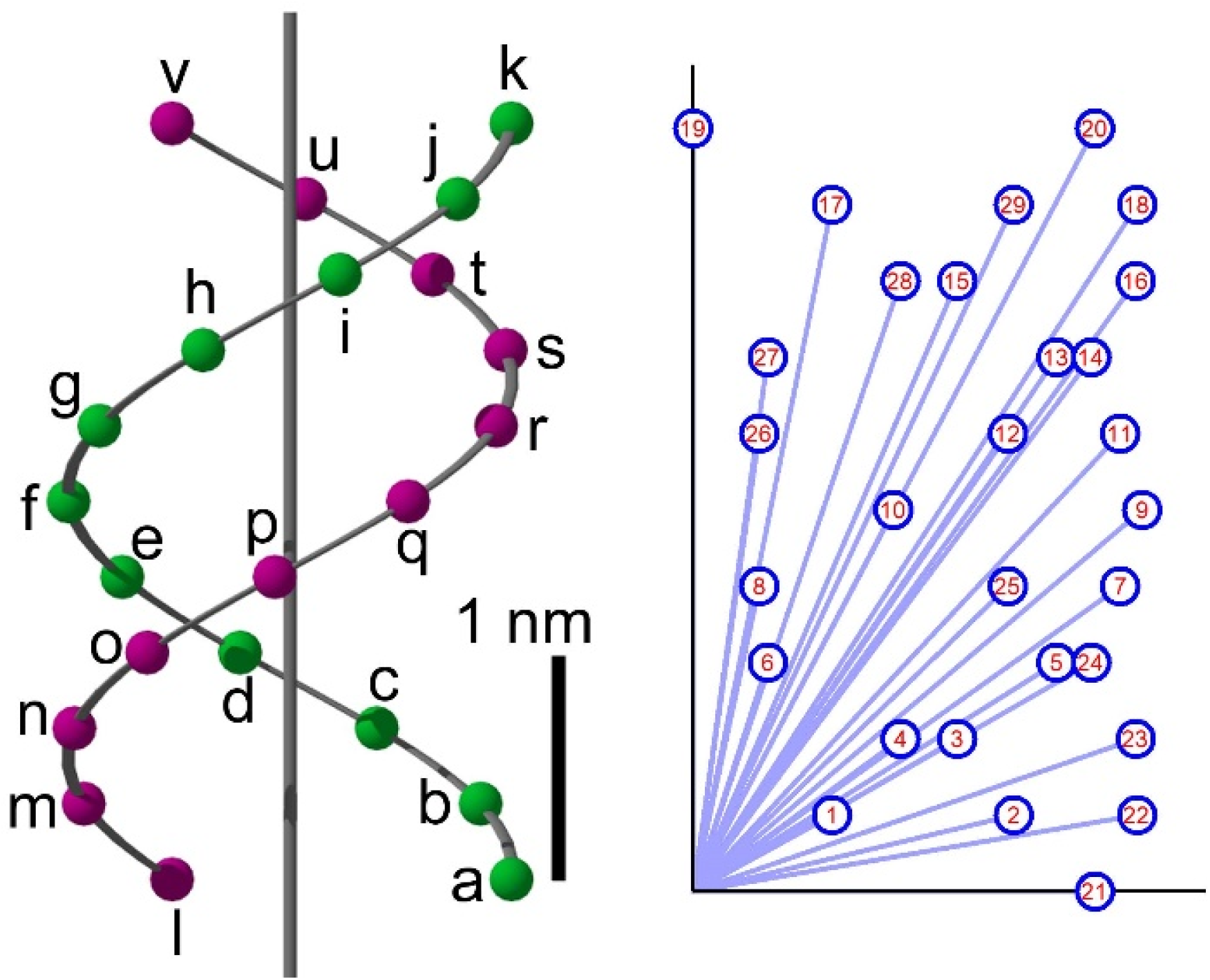
The relationship between the internal vectors in DNA structure (left) and the vectors on the CAP (right). The vectors on the CAP are numbered in red, and the internal vectors represented by each CAP vector are listed below. There are many overlapping vectors, and single number (red) is assigned to them.

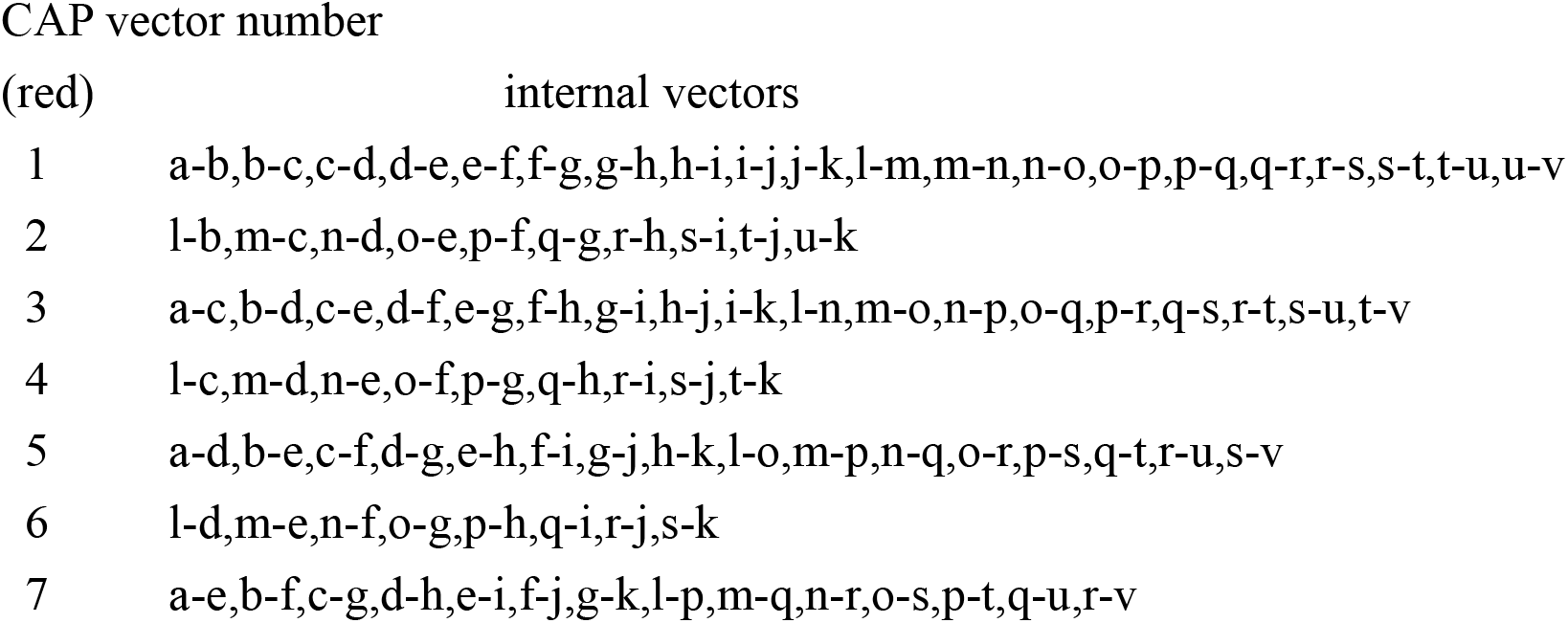
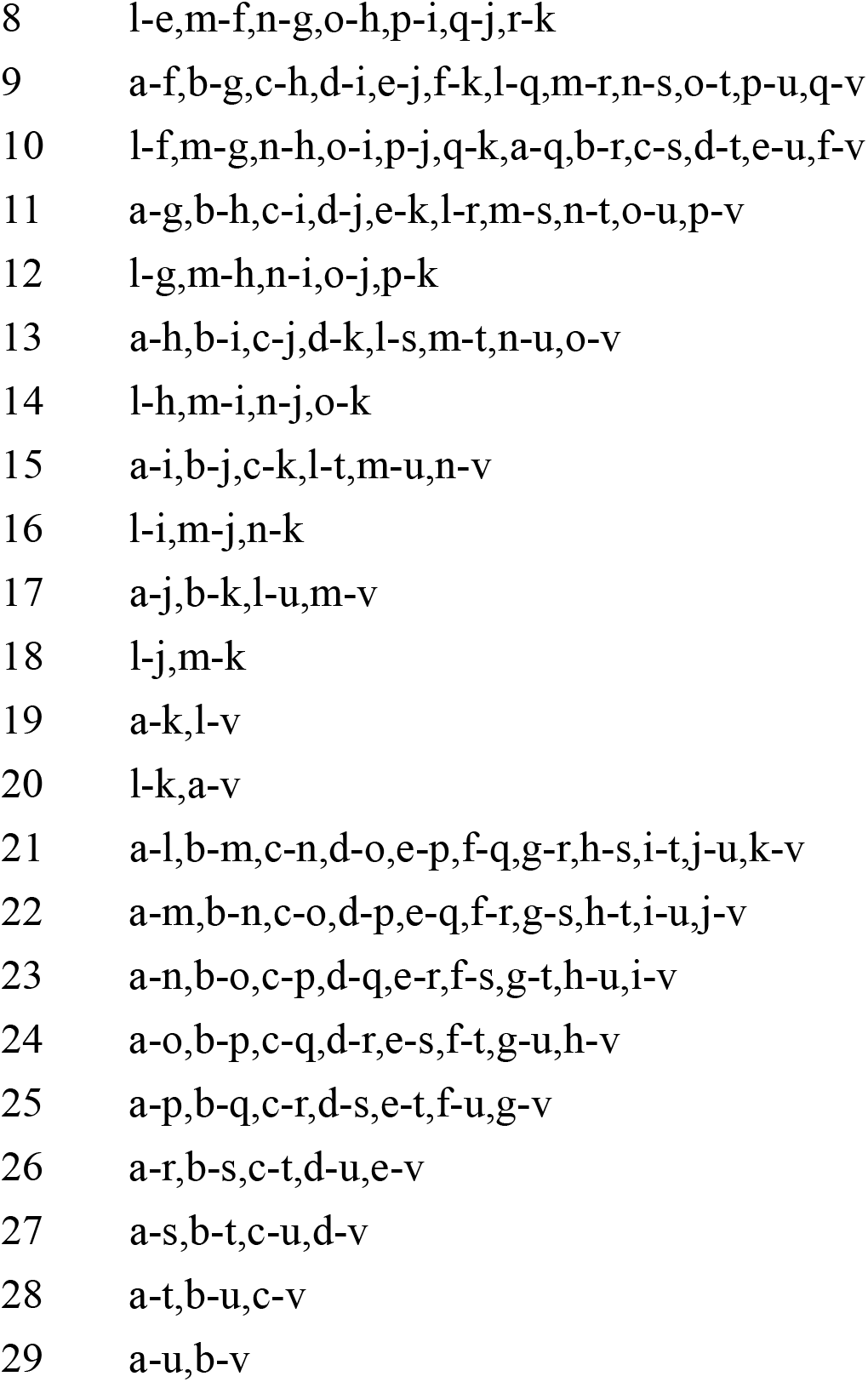

**Supplementary Fig. 3.**
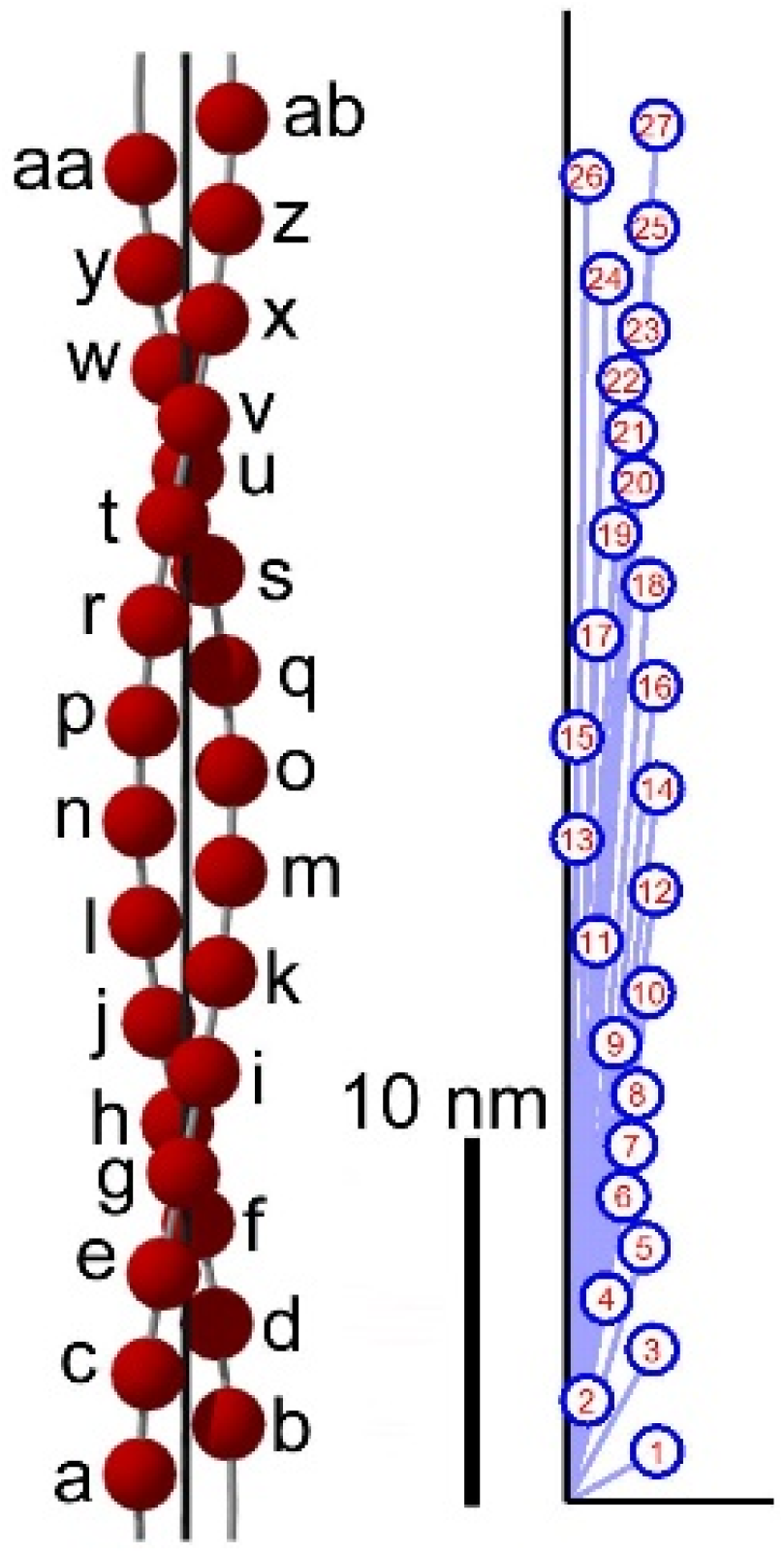
The relationship between the internal vectors in F-actin structure (left) and the vectors on the CAP (right). See legends to Suppl. Fig. 1 for the way of representation.

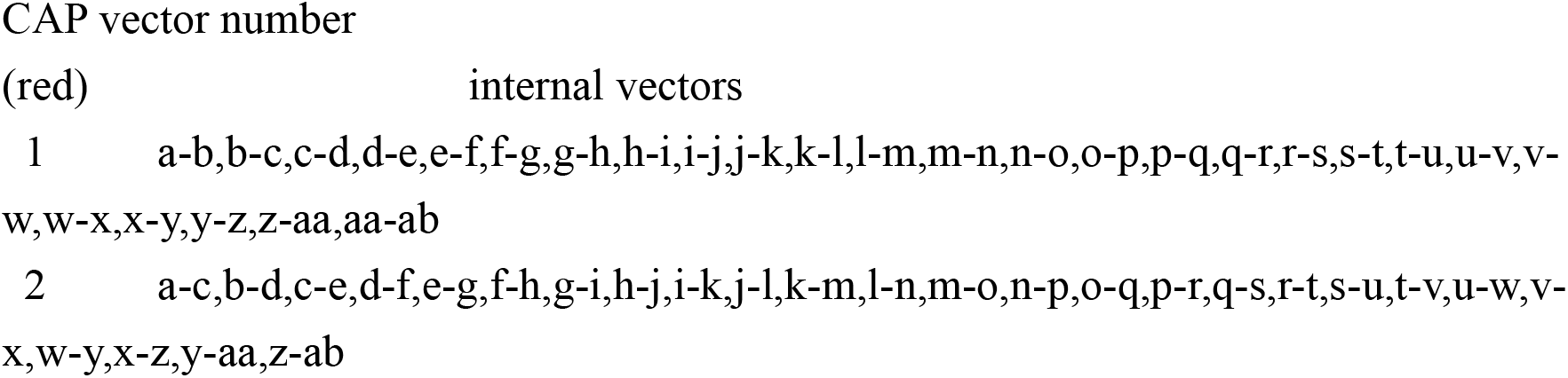
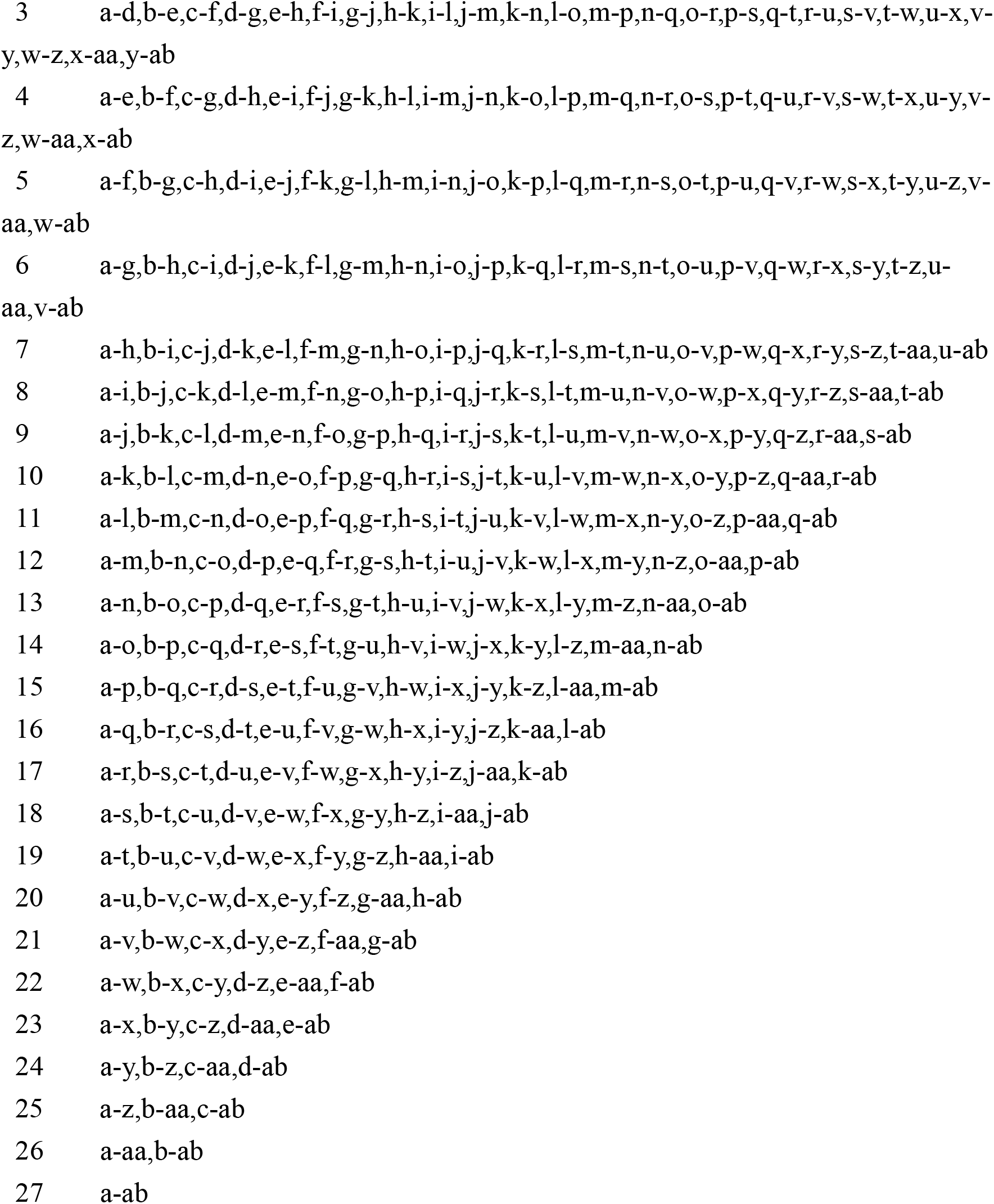

### Legends to supplementary movies

**Movie S1.**

A 3-D object consisting of 3 points, located at the vertices of an equilateral triangle. From its cylindrically averaged Patterson function (CAP), its 3-D structure is restored as demonstrated in Movie S2.

**Movie S2.**

Procedure of restoration of non-rotationally averaged 3-D structure of the object in Movie S1 from its CAP.

**Movie S3.**

Original structure of a double-stranded DNA strand, 11 base pairs (left, phosphate positions only) and the structure restored from its CAP by using the same procedure as in Movie S2 (right).

**Movie S4.**

Original structure of a 28/13 helix of actin, 28 monomers (left) and the structure restored from its CAP by using the same procedure as in Movie S2 (right).

**Movie S5.**

Structure of the myosin filament of bumblebee flight muscle, solved from the CAP calculated from the diffraction pattern from actin-extracted muscle fibers. Four helical strands of myosin heads are colored differently.

